# Can expectation suppression be explained by reduced attention to predictable stimuli?

**DOI:** 10.1101/2020.06.19.154427

**Authors:** Arjen Alink, Helen Blank

## Abstract

The expectation-suppression effect – reduced stimulus-evoked responses to expected stimuli – is widely considered to be an empirical hallmark of reduced prediction errors in the framework of predictive coding. Here we challenge this notion by proposing that that expectation suppression could be explained by a reduced attention effect. Specifically, we argue that reduced responses to predictable stimuli can also be explained by a reduced saliency-driven allocation of attention. We base our discussion mainly on findings in the visual cortex and propose that resolving this controversy requires the assessment of qualitative differences between the ways in which attention and ‘surprise’ enhance brain responses.

## Introduction

Over the past decade, predictive coding theories have gained significant traction with high-profile findings across a range of sensory and cognitive domains (Friston, 2005; Rao & Ballard, 1999; Summerfield & de Lange, 2014; Walsh et al., 2020). Central to these theories lies the idea that the computation of prediction errors forms a principal computational mechanism in the brain. One phenomenon that has been frequently reported and cited as empirical support for such theories is the expectation-suppression phenomenon - reduced stimulus-evoked responses to expected stimuli (Alink et al., 2010; Blank & Davis, 2016; Richter et al., 2018; Todorovic & de Lange, 2012; Walsh & McGovern, 2018). The exact neural mechanism via which predictive coding could give rise to expectation suppression, however, remains unclear. For example, a debate is still ongoing on whether it results from dampening (or explaining away) of neural representations for predictable sensory input or from more selective (or ‘sharpened’) neural representation of predictable stimuli (Friston, 2005; Kok et al., 2012; Richter et al., 2018; Walsh & McGovern, 2018). Furthermore, it has been argued that previously reported expectation-suppression effects can also be attributed to enhanced responses to surprising events (Feuerriegel et al., 2020).

In this perspective, we expand the ongoing discussion about the neural mechanism underlying expectation-suppression by suggesting that expectation suppression could result from reduced attention to predictable stimuli (Aitchison & Lengyel, 2017). In short, we argue that surprising stimuli are more salient and therefore more likely to attract attention (Desimone & Duncan, 1995; Itti & Baldi, 2009). We base our discussion primarily on neuroscientific findings from the visual domain, but this argument should apply beyond visual cortex to other domains in which expectation suppression has been demonstrated (such as basic auditory (Todorovic & de Lange, 2012) or speech processing (Blank & Davis, 2016)).

### Two alternative explanations for expectation suppression

A prominent view suggests that expectation suppression may occur irrespectively of whether stimuli are attended (Kok et al., 2012; Summerfield & Egner, 2009, 2016). One argument for this notion was that better detection of predictable stimuli paired with reduced stimulus evoked responses is difficult to reconcile with the positive neural gain mechanism associated with attention (Alink et al., 2010; Treue & Trujillo, 1999).

This independency between attention and expectation has been challenged by a recent study by Richter and de Lange (2019) which shows that diverting attention away from a predictable image completely abolishes the expectation-suppression effect in the visual cortex. Specifically, conditional probabilities of presented object images no longer had an effect on the amplitude of visual cortex responses if the participants’ attention was drawn away from the object images by a central fixation task. Nonetheless, this finding is interpreted in light of a ***predictive-coding based explanation*** (PBE) by proposing that attending an object boosts prediction error coding. Specifically, Richter et al (2019) suggest that enhanced prediction error coding results from attention increasing the gain of prediction error units (Feldman & Friston, 2010; Friston, 2009; Spratling, 2008) or from attention being a prerequisite for the generation of predictions. Note that, in contrast to this attention-driven boost of error coding, attention has also been reported to suppress prediction error coding in auditory-scene analysis (Sohoglu & Chait, 2016).

In this perspective, we want to discuss an alternative ***attention-based explanation*** (ABE) of expectation suppression which attributes reduced responses to predictable stimuli to a reduced allocation of attentional resources. This alternative is motivated by the notion that attention results from a competition between stimulus representations for the limited information processing capacity of the brain (Desimone & Duncan, 1995). This competition is biased towards behaviourally relevant stimuli, e.g., via enhanced responses to task-relevant and salient stimuli. According to this definition, saliency is a stimulus characteristic that automatically results from deviations from both its physical context and from ‘higher-level’ context based on natural image statistics and experience (Desimone & Duncan, 1995; Hickey & Peelen, 2015; Kanan et al., 2009). An important implication of this biased competition model is that attention towards a stimulus is not only determined top-down by its task-relevance, but also by its bottom-up saliency. These two forms of attention have also been differentiated as endogenous and exogenous attention, respectively (Chun & Wolfe, 2001). While the former is a voluntary system, the latter is an involuntary system that corresponds to an automatic orienting response to unexpected stimulation (Carrasco, 2011). To illustrate, the sound of a fire-alarm will capture your full attention no matter how strictly you have been instructed to perform a visual discrimination task. Accordingly, ABE proposes that enhanced responses to unexpected stimuli in sensory brain areas reflect a greater saliency-driven allocation of attentional resources towards these stimuli (Figure 1).

**Figure 1.**
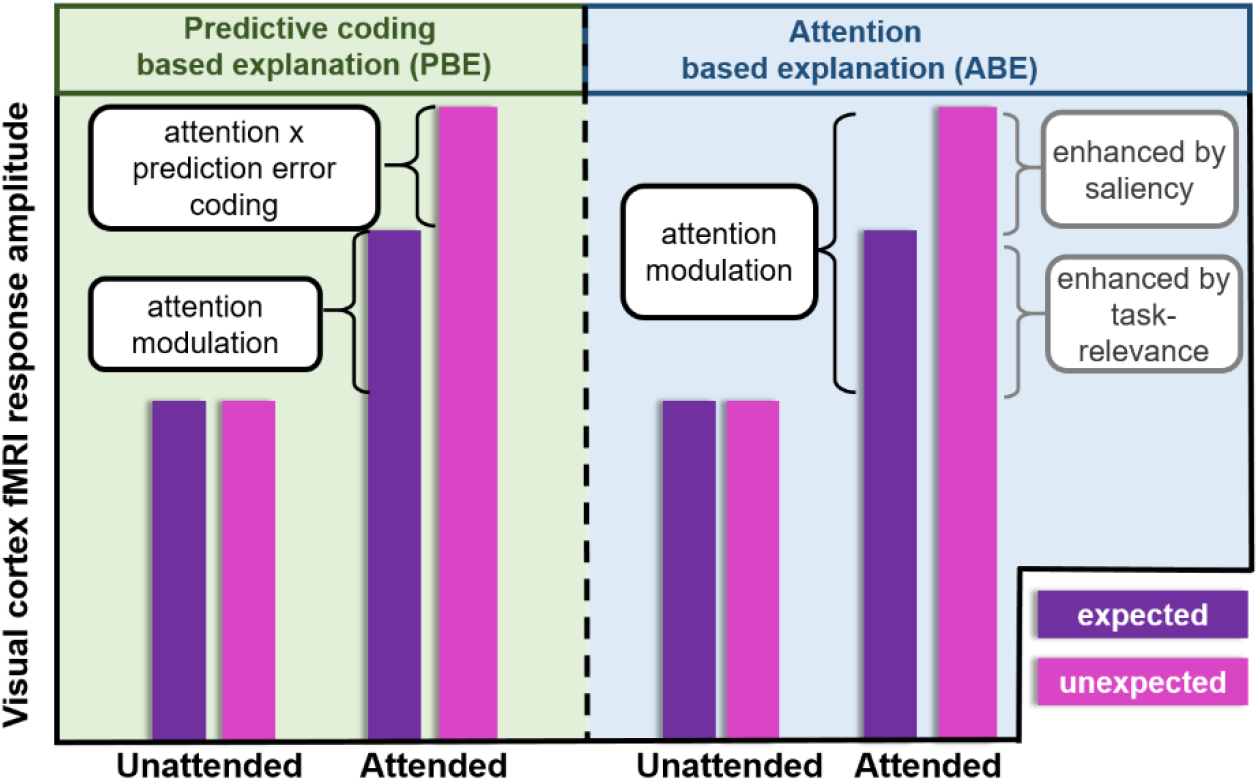
Predictive-coding based explanation. (PBE, left) attention increases the responsiveness of neurons sensitive to the discrepancy of the sensory input and the expected stimulus. suppression as resulting from a reduced stimulus. **Attention-based explanation** (ABE, right) explains expectation saliency of expected stimuli.

The ABE does not equate stimulus probability to attention. Instead, it stresses that one determinant of attention allocation, stimulus saliency, depends on the extent to which stimuli violate the regularities of their spatial or temporal context. For example, a stimulus is more salient when it is the deviant stimulus in an oddball paradigm as compared to when it is part of a random stimulus sequence. Note that, in contrast to the PBE, the ABE does not strictly require prediction-error coding as the expectation suppression effect is fully explained by reduced stimulus saliency (however, see Spratling, 2008). Finally, we would like to point out that the ABE does not preclude the possibility that attention is drawn towards predictable events under certain circumstances (e.g., when predictable stimuli are task-relevant, Alamia & Zénon, 2016).

For illustrative purposes, let us evaluate whether the ABE can accommodate the finding by Richter and de Lange (2019) that expectation suppression only occurs in the visual cortex when images are attended. As outlined above, unexpected stimuli can be thought of as more capable of capturing attention due to their saliency, which is compatible with enhanced pupil dilation for unexpected stimuli reported in this study (Richter & de Lange, 2019; van der Brink et al., 2016). Therefore, predictable images should have received less attention which could explain the reduced associated responses in the visual cortex. The reported absence of this expectation-suppression effect for the central fixation task also fits with the ABE as this task is argued to reduce attention allocation towards expected and unexpected images (Richter & de Lange, 2019).

### Why it is important to differentiate between a predictive coding and an attention-based explanation of expectation suppression?

Over the last decades, theories of predictive coding became more and more popular and expectation suppression is an observation that has frequently been interpreted as supporting such theories. Although we agree that reduced prediction error coding provides an appealing explanation for reduced responses to expected stimuli, we here want to join others in requesting that predictive coding should not be accepted as an ‘a-priori truth’ and that we should remain open to considering alternative explanations to prevent this theoretical framework from turning into a ‘just-so story’ that is able to accommodate any empirical finding (Bowers & Davis, 2012; Cao, 2020; Heilbron & Chait, 2017; Walsh et al., 2020). Note, however, that the same issue applies to the concept of attention, which appears to be equally ill-defined (Hommel et al., 2019).

In addition, differentiating between the two alternative explanations of expectation suppression is important because they make different predictions about what the increased neural activity for unexpected stimuli signifies: The PBE suggests that this increased activity encodes the discrepancy between the actual and expected input to a brain area, while the ABE suggests that this enhanced activity results from increased attention allocation towards salient stimuli. As pointed out by Spratling (2008), it is possible that stimulus saliency is computed in a similar manner as prediction errors. However, even if stimulus saliency can be equated to a prediction error coding process, the ABE and the PBE differ with regard to the order of attention effects and the encoding of prediction errors/stimulus saliency: The PBE proposes that attention facilitates prediction error coding (Feldman & Friston, 2010; Friston, 2009; Richter & de Lange, 2019) while the ABE suggests that attention effects result from stimulus saliency (Desimone & Duncan, 1995, Kanan et al., 2009). Therefore, the critical difference appears to be that the PBE suggests that attention is *a cause* of enhanced prediction-error coding, while the ABE implies that attention is *an effect* of stimulus saliency.

Since the ABE and the PBE can both explain expectation suppression, one would ideally want to apply Ockham’s razor to determine which provides the more parsimonious explanation (Heilbron & Chait, 2017). Such an approach, however, is challenging because attention and predictive coding models are typically not formulated at the same level of description (Marr, 1982). Therefore, resolving whether the ABE or the PBE is more parsimonious requires a reformulation of these explanations at a comparable level of description (e.g., see Spratling, 2008). This would help shed light on whether predictive coding models, whose complexity is increasing (Press et al., 2020), offer the most parsimonious explanation of the expectation-suppression effect.

### The challenge of orthogonalizing attention and expectation effects in the human brain

As pointed out by Summerfield and Egner (Summerfield & Egner, 2016), insights into how expectation affects sensory processing in the human brain requires experiments that explicitly orthogonalize expectation and attention effects. Specifically, they suggest a normative distinction between feature-based attention (FBA) and feature-based expectation (FBE), where FBA is manipulated by task-relevance, while FBE is manipulated by stimulus probability. Importantly, FBA is proposed to enhance sensitivity to attended features, while FBE is proposed to bias perception towards the expected stimulus by adjusting decision criteria. This definition of FBA restricts attention effects to those driven by task-relevance. However, as we pointed out above, context-violating (and therefore salient) stimuli can capture exogenous attention irrespective of the task at hand. Resolving this issue, requires neuroimaging paradigms specifically geared towards determining if the neural consequence of surprise can be dissociated from endogenous and exogenous attention effects that are not related to stimulus probability (as outlined below).

In addition, we would like to highlight that it is unclear whether expectation and attention effects map onto brain mechanisms that are computationally or neurophysiologically dissociable (Summerfield & de Lange, 2014). For example, the question arises whether expectation suppression and expectation-induced sharpening of sensory representations in early sensory brain areas are consistent with the implementation of FBE, because perceptual biases are typically thought to result from activity in higher level areas (Alink et al., 2010; Heekeren et al., 2008; Kok et al., 2012; Rungratsameetaweemana & Serences, 2019).

In conclusion, it is important to acknowledge the possibility that unpredictable stimuli capture more exogenous attention and to address this challenge by differentiating neurophysiological consequences of expectation and attention.

### Possible approaches for testing whether expectation suppression is a pure attention effect

How could we determine whether expectation suppression results from predictive coding or from reduced attention to expected stimuli? As pointed out above (e.g., Figure 1), univariate responses cannot differentiate between these two possibilities, because PBE and ABE both predict reduced responses to expected stimuli (Alink et al., 2010; Blank & Davis, 2016). Also, the observation that stimulus predictability improves perceptual decisions (Kok et al., 2012; Schwiedrzik et al., 2007) does not enable us to rule out PBE or ABE, because they both are consistent with perceptual decisions being made based on sensory input *and* prior expectations - which in general leads to improved perceptual decisions for predictable stimuli.

#### Designs controlling endogenous and exogenous attention

One way to strengthen the case for PBE would be to demonstrate qualitative differences between *how* ‘surprise’ and a ‘pure top-down attention effect’ enhance brain responses. This is possible because, although stimulus probabilities are likely to affect exogenous attention (via stimulus saliency), endogenous task-driven attention can be manipulated independently from stimulus probability (Summerfield & de Lange, 2014). This requires experiments that independently manipulate the probability and task relevance of stimuli to directly assess qualitative differences between surprise-related brain response enhancement and a pure endogenous attentional gain effect. The demonstration of such a difference would speak against the ABE, at least when assuming that saliency and task relevance induce the same attentional neural gain effect (Desimone & Duncan, 1995). Previous work using such an approach of orthogonalizing task-driven, i.e., endogenous attention and stimulus probability in the visual (Kok, et al., 2012; Marzecová et al., 2017, 2018) and auditory (Saupe et al., 2013; Timm et al., 2013) domain revealed overlapping and interacting but importantly also independent modulations of these manipulations.

Next, the ABE of expectation suppression could be falsified by orthogonally varying stimulus probability and the degree to which stimuli attract exogenous attention. For example, one could present images with varying saliency (e.g., via different levels of emotion (Carretié, 2014)), while independently rendering images expected and unexpected (e.g., via statistical learning (Richter et al., 2019)). A demonstration of qualitatively distinct effects of stimulus predictability and saliency on brain responses would provide evidence against surprise and exogenous attention affecting brain responses via a common neural mechanism.

#### Multivariate decoding approaches

Alternatively, one could falsify the ABE by conclusively demonstrating that expectation suppression enhances stimulus decodability based on the stimulus evoked neural responses (e.g., based on fMRI response patterns). Attention is known to enhance the responses of neurons tuned to the attended stimulus, which has been shown to give rise to greater fMRI-pattern based stimulus decodability (Kamitani & Tong, 2005). Therefore, according to the ABE, unexpected stimuli should be more decodable. In contrast, it is unclear whether according to the PBE unexpected or expected stimuli should be more decodable. If reduced responses to expected stimuli result from ‘explaining away’ of predicted sensory input (Clark, 2013), unexpected stimuli should be more decodable. However, stimulus predictability has been associated with enhanced fMRI pattern decodability which has led to the proposal that expectations sharpen neural representations of expected stimuli (Kok et al., 2012). Such expectation-induced sharpening would contradict the ABE. At present, however, expectation-driven sharpening remains controversial because enhanced decodability has also been observed for unexpected stimuli (Jiang et al., 2013; Kumar et al., 2017; Richter et al., 2018; Tang et al., 2018) and given the observation that a conjunction of reduced fMRI responses and enhanced decodability can also be explained by a local-scaling effect (Alink et al., 2018). In conclusion, more evidence for sharpened responses to expected stimuli is needed to rule out the ABE.

#### Methods with high laminar precision (e.g., 7T fMRI)

According to the theoretical framework of predictive coding, dampened responses to expected stimuli result from attenuated responses in ‘prediction error units’ and sharpened responses in ‘hypothesis units’ (Friston, 2005), which are proposed to reside in different cortical layers (de Lange et al., 2018; Kok et al., 2016; Muckli et al., 2015). Therefore, methods with high laminar precision (e.g., 7T fMRI) should be able to validate PBE by establishing a link between expectation suppression and laminar-specific dampening and sharpening effects.

#### Methods with high temporal resolution

Finally, one could discriminate between the ABE and the PBE by using temporally resolved measurements (e.g., using EEG or MEG), because the explanations differ with regard to the relative timing of attention allocation and prediction error coding/stimulus saliency encoding.

The ABE explains increased responses to unexpected stimuli as enhanced exogenous attention allocation towards these unexpected stimuli. Previously, it has been demonstrated that exogenous attention effects precede endogenous attention effects (Hickey et al., 2010). Therefore, one could test the validity of the ABE by demonstrating that expectation effects precede endogenous attention effects, for example, by measuring the onset of lateralized effects of expectation and task-driven attention with EEG (Hickey et al., 2010; Luck & Hillyard, 1994). In contrast, the PBE suggests that neural effects of endogenous attention allocation precede suppressive effects of expectation (Richter et al, 2019) and that predictive coding takes place during later stages of sensory processing (Press et al., 2020).

Alternatively, on could determine whether enhanced responses to unexpected stimuli result from exogenous attention by testing if they are associated with a selective enhancement of the N1 component, which has been shown to index exogenously driven attention shifts (Natale et al., 2006).

### Applicability of ABE to other expectation effects

Here, we discuss whether the expectation suppression effect can be equally well explained by attention as by predictive coding. It will be of interest to explore if this also applies to other modulations of brain responses typically associated with predictive coding, such as omission-evoked responses resembling responses to the expected stimulus (Bendixen et al., 2009) and feature-specific pre-stimulus activity induced by expectations (Fiser et al., 2016; Kok et al., 2017). ABE, at least partially, appears to be able to account for these phenomena because 1) it explains enhanced responses to unexpected omissions as an effect of greater saliency-driven attention and 2) because ABE also requires neural representations of spatiotemporal context (Desimone & Duncan, 1995). However, ABE does not provide a straight-forward explanation for why omission-induced responses should encode the features of unexpected stimuli (Bendixen et al., 2009). Therefore, progress in disentangling predictive coding effects from attention effects could be made by assessing the explanatory power of attention and predictive coding theories across a wide range of neurophysiological effects of expectation.

### Predictive coding, moving beyond “New Labels for Old Ideas”

A current debate around theories of predictive coding is whether they provide new constraints on brain function that have not already been provided by traditional models (Cao, 2020). In this perspective we contribute to this debate by discussing the possibility that expectation suppression, an empirical finding frequently interpreted as support for predictive coding theories, can be accounted for by traditional models of attention. In short, we argue that enhanced brain responses to surprising stimuli can be accounted for by an enhanced allocation of exogenous attention. In this commentary, we discuss how this issue can be resolved with experiments that specifically assess the different ways in which expectation and attention affect brain responses.

## Acknowledgements

We thank Rik Henson, David Richter, and Floris de Lange for thoughtful comments on earlier versions of this commentary.

## References

Aitchison, L., & Lengyel, M. (2017). With or without you: Predictive coding and Bayesian inference in the brain. SI: 46: Computational Neuroscience (2017), 46(Supplement C), 219–227. https://doi.org/10.1016/j.conb.2017.08.010

Alamia, A., & Zénon, A. (2016). Statistical Regularities Attract Attention when Task-Relevant. Frontiers in Human Neuroscience, 10. https://doi.org/10.3389/fnhum.2016.00042

Alink, A., Schwiedrzik, C. M., Kohler, A., Singer, W., & Muckli, L. (2010). Stimulus predictability reduces responses in primary visual cortex. The Journal of Neuroscience, 30(8), 2960–2966. https://doi.org/10.1523/JNEUROSCI.3730-10.2010

Bendixen, A., Schröger, E., & Winkler, I. (2009). I Heard That Coming: Event-Related Potential Evidence for Stimulus-Driven Prediction in the Auditory System. The Journal of Neuroscience, 29(26), 8447. https://doi.org/10.1523/JNEUROSCI.1493-09.2009

Blank, H., & Davis, M. H. (2016). Prediction Errors but Not Sharpened Signals Simulate Multivoxel fMRI Patterns during Speech Perception. PLOS Biology, 14(11), e1002577. https://doi.org/10.1371/journal.pbio.1002577

Bowers, J. S., & Davis, C. J. (2012). Bayesian just-so stories in psychology and neuroscience. Psychological Bulletin, 138(3), 389–414. https://doi.org/10.1037/a0026450

Cao, R. (2020). New Labels for Old Ideas: Predictive Processing and the Interpretation of Neural Signals. Review of Philosophy and Psychology. https://doi.org/10.1007/s13164-020-00481-x

Carrasco, M. (2011). Visual attention: The past 25 years. Vision Research, 51(13), 1484–1525. https://doi.org/10.1016/j.visres.2011.04.012

Carretié, L. (2014). Exogenous (automatic) attention to emotional stimuli: A review. Cognitive, Affective, & Behavioral Neuroscience, 14(4), 1228–1258. https://doi.org/10.3758/s13415-014-0270-2

Chun, M., & Wolfe, J. (2001). Visual Attention. In E. B. Goldstein (Ed.), Blackwell Handbook of Perception (pp. 2–335). Blackwell.

de Lange, F. P., Heilbron, M., & Kok, P. (2018). How Do Expectations Shape Perception? Trends in Cognitive Sciences. https://doi.org/10.1016/j.tics.2018.06.002

den Ouden, H. E. M., Friston, K. J., Daw, N. D., McIntosh, A. R., & Stephan, K. E. (2009). A Dual Role for Prediction Error in Associative Learning. Cerebral Cortex, 19(5), 1175–1185. https://doi.org/10.1093/cercor/bhn161

Desimone, R., & Duncan, J. (1995). Neural Mechanisms of Selective Visual Attention. Annu Rev Neurosci, 30.

Feldman, H., & Friston, K. J. (2010). Attention, uncertainty, and free-energy. Frontiers in Human Neuroscience, 4, 215. https://doi.org/10.3389/fnhum.2010.00215

Feuerriegel, D., Vogels, R., & Kovács, G. (2020). Evaluating the Evidence for Expectation Suppression in the Visual System. PsyArXiv. https://doi.org/10.31234/osf.io/y7639

Fiser, A., Mahringer, D., Oyibo, H. K., Petersen, A. V., Leinweber, M., & Keller, G. B. (2016). Experience-dependent spatial expectations in mouse visual cortex. Nature Neuroscience, 19(12), 1658–1664. https://doi.org/10.1038/nn.4385

Friston, K. (2005). A theory of cortical responses. Philos Trans R Soc London [Biol], 360(1456), 815–836.

Friston, Karl. (2009). The free-energy principle: A rough guide to the brain? Trends in Cognitive Sciences, 13(7), 293–301.

Heekeren, H. R., Marrett, S., & Ungerleider, L. G. (2008). The neural systems that mediate human perceptual decision making. Nature Reviews Neuroscience, 9(6), 467–479. https://doi.org/10.1038/nrn2374

Heilbron, M., & Chait, M. (2017). Great expectations: Is there evidence for predictive coding in auditory cortex? Neuroscience. https://doi.org/10.1016/j.neuroscience.2017.07.061

Hickey, C., & Peelen, M. V. (2015). Neural Mechanisms of Incentive Salience in Naturalistic Human Vision. Neuron, 85(3), 512–518. https://doi.org/10.1016/j.neuron.2014.12.049

Hickey, C., van Zoest, W., & Theeuwes, J. (2010). The time course of exogenous and endogenous control of covert attention. Experimental Brain Research. Experimentelle Hirnforschung. Experimentation Cerebrale, 201(4), 789–796. https://doi.org/10.1007/s00221-009-2094-9

Hommel, B., Chapman, C. S., Cisek, P., Neyedli, H. F., Song, J.-H., & Welsh, T. N. (2019). No one knows what attention is. Attention, Perception, & Psychophysics, 81(7), 2288–2303. https://doi.org/10.3758/s13414-019-01846-w

Itti, L., & Baldi, P. (2009). Bayesian surprise attracts human attention. Visual Attention: Psychophysics, Electrophysiology and Neuroimaging, 49(10), 1295–1306. https://doi.org/10.1016/j.visres.2008.09.007

Kanan, C., Tong, M. H., Zhang, L., & Cottrell, G. W. (2009). SUN: Top-down saliency using natural statistics. Visual Cognition, 17(6-7), 979–1003. https://doi.org/10.1080/13506280902771138

Kok, P., Bains, L. J., van Mourik, T., Norris, D. G., & de Lange, F. P. (2016). Selective Activation of the Deep Layers of the Human Primary Visual Cortex by Top-Down Feedback. Current Biology, 26(3), 371–376. https://doi.org/10.1016/j.cub.2015.12.038

Kok, P., Jehee, J. F. M., & de Lange, F. P. (2012). Less Is More: Expectation Sharpens Representations in the Primary Visual Cortex. Neuron, 75(2), 265–270. https://doi.org/10.1016/j.neuron.2012.04.034

Kok, P., Rahnev, D., Jehee, J. F., Lau, H. C., & de Lange, F. P. (2012). Attention reverses the effect of prediction in silencing sensory signals. Cereb Cortex, 22(9), 2197–2206. https://doi.org/10.1093/cercor/bhr310

Kok, Peter, Mostert, P., & de Lange, F. P. (2017). Prior expectations induce prestimulus sensory templates. Proceedings of the National Academy of Sciences. https://doi.org/10.1073/pnas.1705652114

Luck, S. J., & Hillyard, S. A. (1994). Electrophysiological correlates of feature analysis during visual search. Psychophysiology, 31(3), 291–308. https://doi.org/10.1111/j.1469-8986.1994.tb02218.x

Marzecová, A., Schettino, A., Widmann, A., SanMiguel, I., Kotz, S. A., & Schröger, E. (2018). Attentional gain is modulated by probabilistic feature expectations in a spatial cueing task: ERP evidence. Scientific Reports, 8(1), 54. https://doi.org/10.1038/s41598-017-18347-1

Marzecová, A., Widmann, A., SanMiguel, I., Kotz, S. A., & Schröger, E. (2017). Interrelation of attention and prediction in visual processing: Effects of task-relevance and stimulus probability. Biological Psychology, 125, 76–90. https://doi.org/10.1016/j.biopsycho.2017.02.009

Muckli, L., De Martino, F., Vizioli, L., Petro, L. S., Smith, F. W., Ugurbil, K., Goebel, R., & Yacoub, E. (2015). Contextual Feedback to Superficial Layers of V1. Current Biology, 25(20), 2690–2695. https://doi.org/10.1016/j.cub.2015.08.057

Natale, E., Marzi, C. A., Girelli, M., Pavone, E. F., & Pollmann, S. (2006). ERP and fMRI correlates of endogenous and exogenous focusing of visual-spatial attention. European Journal of Neuroscience, 23(9), 2511–2521. https://doi.org/10.1111/j.1460-9568.2006.04756.x

Press, C., Kok, P., & Yon, D. (2020). The Perceptual Prediction Paradox. Trends in Cognitive Sciences, 24(1), 13–24. https://doi.org/10.1016/j.tics.2019.11.003

Rao, R. P., & Ballard, D. H. (1999). Predictive coding in the visual cortex: A functional interpretation of some extra-classical receptive-field effects. Nat Neurosci, 2(1), 79–87. https://doi.org/10.1038/4580

Richter, D., & de Lange, F. P. (2019). Statistical learning attenuates visual activity only for attended stimuli. ELife, 8, e47869. https://doi.org/10.7554/eLife.47869

Richter, D., Ekman, M., & de Lange, F. P. (2018). Suppressed sensory response to predictable object stimuli throughout the ventral visual stream. The Journal of Neuroscience. https://doi.org/10.1523/JNEUROSCI.3421-17.2018

Rungratsameetaweemana, N., & Serences, J. T. (2019). Dissociating the impact of attention and expectation on early sensory processing. Current Opinion in Psychology, 29, 181–186. https://doi.org/10.1016/j.copsyc.2019.03.014

Saupe, K., Widmann, A., Trujillo-Barreto, N. J., & Schröger, E. (2013). Sensorial suppression of self-generated sounds and its dependence on attention. International Journal of Psychophysiology, 90(3), 300–310. https://doi.org/10.1016/j.ijpsycho.2013.09.006

Schwiedrzik, C. M., Alink, A., Kohler, A., Singer, W., & Muckli, L. (2007). A spatiotemporal interaction on the apparent motion trace. Vision Research, 47(28), 3424–3433.

Sohoglu, E., & Chait, M. (2016). Detecting and representing predictable structure during auditory scene analysis. ELife, 5, e19113. https://doi.org/10.7554/eLife.19113

Spratling, M. W. (2008). Reconciling predictive coding and biased competition models of cortical function. Front Comput Neurosci, 2, 4. https://doi.org/10.3389/neuro.10.004.2008

Summerfield, C., & de Lange, F. P. (2014). Expectation in perceptual decision making: Neural and computational mechanisms. Nat Rev Neurosci, 15, 745–756. https://doi.org/10.1038/nrn3838

Summerfield, Christopher, & Egner, T. (2009). Expectation (and attention) in visual cognition. Trends in Cognitive Sciences, 13(9), 403–409.

Summerfield, Christopher, & Egner, T. (2016). Feature-based attention and featurebased expectation. Trends in Cognitive Sciences, 20(6), 401–404.

Timm, J., SanMiguel, I., Saupe, K., & Schröger, E. (2013). The N1-suppression effect for self-initiated sounds is independent of attention. BMC Neuroscience, 14(1), 2. https://doi.org/10.1186/1471-2202-14-2

Todorovic, A., & de Lange, F. P. (2012). Repetition Suppression and Expectation Suppression Are Dissociable in Time in Early Auditory Evoked Fields. Journal of Neuroscience, 32(39), 13389–13395. https://doi.org/10.1523/jneurosci.2227-12.2012

Treue, S., & Trujillo, J. C. M. (1999). Feature-based attention influences motion processing gain in macaque visual cortex. Nature, 399(6736), 575–579. https://doi.org/10.1038/21176

van der Brink, R. L. van den, Murphy, P. R., & Nieuwenhuis, S. (2016). Pupil Diameter Tracks Lapses of Attention. PLOS ONE, 11(10), e0165274. https://doi.org/10.1371/journal.pone.0165274

Walsh, K. S., & McGovern, D. P. (2018). Expectation Suppression Dampens Sensory Representations of Predicted Stimuli. The Journal of Neuroscience, 38(50), 10592. https://doi.org/10.1523/JNEUROSCI.2133-18.2018

Walsh, K. S., McGovern, D. P., Clark, A., & O’Connell, R. G. (2020). Evaluating the neurophysiological evidence for predictive processing as a model of perception. Annals of the New York Academy of Sciences, 1464(1), 242–268. https://doi.org/10.1111/nyas.14321

